# Misalignment of selection, plasticity, and among-individual variation: A test of theoretical predictions with *Peromyscus maniculatus*

**DOI:** 10.1101/2020.12.07.415190

**Authors:** Monica Anderson Berdal, Ned A. Dochtermann

**Author notes:** Author contributions: M.A.B and N.A.D. conceived the project and supervised the gathering of data; M.A.B. supervised and conducted behavioural trials, analysed the data, and wrote the first draft of the manuscript. N.A.D., and M.A.B. wrote the finished manuscript.

## Abstract

Genetic variation and phenotypic plasticity are predicted to align with selection surfaces, a prediction that has rarely been empirically tested. Understanding the relationship between sources of phenotypic variation, i.e. genetic variation and plasticity, with selection surfaces improves our ability to predict a population’s ability to adapt to a changing environment and our understanding of how selection has shaped phenotypes. Here, we estimated the (co)variances among three different behaviors (activity, aggression, and anti-predator response) in a natural population of deer mice (*Peromyscus maniculatus*). Using multi-response generalized mixed effects models, we divided the phenotypic covariance matrix into among- and within-individual matrices. The among-individual covariances includes genetic and permanent environmental covariances (e.g. developmental plasticity) and is predicted to align with selection. Simultaneously, we estimated the within-individual (co)variances, which include reversible phenotypic plasticity. To determine whether genetic variation, plasticity and selection align in multivariate space we calculated the dimensions containing the greatest among-individual variation and the dimension in which most plasticity was expressed (i.e. the dominant eigenvector for the among- and within-individual covariance matrices respectively). We estimated selection coefficients based on survival estimates from a mark-recapture model. Alignment between the dominant eigenvectors of behavioural variation and the selection gradient was estimated by calculating the angle between them, with an angle of 0 indicating perfect alignment. The angle between vectors ranged from 68° to 89°, indicating that genetic variation, phenotypic plasticity, and selection are misaligned in this population. This misalignment could be due to the behaviors being close to their fitness optima, which is supported by low evolvabilities, or because of low selection pressure on these behaviors.

## Introduction

Populations under selection should have phenotypic distributions reflecting this selection (1-3). To maximize fitness the majority of phenotypic variation should therefore align with the selection gradient (**β**, 4, 5, 6). Put another way, the axes of most phenotypic variation should be expressed in the same direction as (i.e. be collinear with) selection gradients. Both genetic and environmental variation contribute to a population’s phenotypic variation (**P = G + E**), and both sources of variation can align with selection (2, 5, 7, 8).

Because traits often covary, selection pressure on one trait affects the mean and variance of other traits and vice versa (1, 9, 10). It is therefore necessary to investigate alignment between variation and selection from a multi-trait perspective. In quantitative genetics, the **G**-matrix summarizes genetic variation and covariation (10) and populations are predicted to evolve faster in the direction of most genetic variation (Schluter 1996). This direction, named the line of least resistance by Schluter (11), is the dominant eigenvector (**g_max_**) of **G**. A population where **β** and **g_max_** are aligned will reach a fitness optimum faster than if there was a misalignment (11). Simultaneously, selection is expected to have shaped **G** and several models have shown that **g_max_** will align with the selection surfaces given enough time and if selection pressures are stable (2, 8).

Misalignment between **g_max_** and **β** can be ameliorated by alignment between selection and environmentally induced variation in phenotypes. Put another way, phenotypic plasticity allows an individual to change their expression of a trait based on the environment and potentially increase their fitness (i.e. adaptive phenotypic plasticity, (12). Adaptive phenotypic plasticity is predicted to evolve in more heterogeneous environments to allow populations to move closer towards trait optimum within each environment (13-15). Consequently, adaptive plasticity should align with the selection gradient (16).

According to recent theory, phenotypic plasticity should also align with **g_max_**, because it maintains, and possibly increases, the genetic variation along the axes where trait optimum vary (7). Consistent with this, Noble, Radersma (6) found via meta-analysis a significant alignment between **g_max_** and direction of plasticity in novel environments. Loss of fitness due to a misalignment between **g_max_** and **β** can therefore be reduced by phenotypic plasticity, contingent on costs of being plastic. While genetic adaptation can only occur over several generations, phenotypic plasticity can change within the same generation, and might therefore be the major contributor to alignment between phenotypic variation and selection in new environments (7, 12, 17). However, because individuals do not show the full repertoire of phenotypes present in a population, costs and/or limits to plasticity are expected to frequently be substantial (18-20). As costs to plasticity increase, **g_max_** is predicted to more quickly align with selection, reducing the need for plasticity and its costs (18, 21).

In natural populations **G**, is difficult to estimate because it requires both a pedigree and trait measurements. Consequently, alignment among **G**, plasticity, and selection gradients has rarely been assessed and existing predictions about their interplay have not been tested. Fortunately, the among-individual variance-covariance matrix (**I**-matrix) can be estimated for a single generation by repeatedly sampling individuals from a population. The **I**-matrix consists of the joint effects of **G** and permanent environmental (**PE**) correlations (22), each explaining approximately 50% of repeatable variance in behaviors (23). Moreover, consistent with Cheverud’s conjecture, phenotypic correlations are highly concordant with genetic correlations (24, 25) and among-individual correlations (26). **I** can therefore be used as a proxy for the G-matrix, albeit with caveats. Similarly, plasticity can be difficult to measure in wild population. However, the within-individual variance-covariance matrix (**W**-matrix) contains the temporary environmental (TE) correlations, i.e. correlation among traits due to changes in the environment within the timeframe of trait measurement. This includes adaptive reversible plasticity (27, 28) and **w_max_**, the dominant eigenvector of **W**, is the direction in which most plasticity is expressed. Thus, using among- and within-individual variance-covariance matrices as proxies for genetic variation and phenotypic plasticity respectively allows us to estimate their alignment with each other and the selection surfaces without the need of a pedigree.

Here we examined alignment between selection, among-individual covariances, and phenotypic plasticity in a wild population of deer mice, *Peromyscus maniculatus* by estimating the angles between the selection gradient (**β**) and the dimensions in phenotypic space containing the greatest amount of among- and within-individual behavioral variation. If fitness optima are stable both within and across generations for our population of deer mice, both among- and within-individual variation are expected to be aligned with the selection gradient and each other.

## Methods

### Study species

A wild population of deer mice was sampled at Cassel wood, Minnesota, USA (Figure S1). Deer mice are highly suited for our questions because repeatability (29, 30) and appreciable additive genetic variation (31) has been previously demonstrated for behaviors similar to those measured here, making among-individual correlations more likely to be a suitable proxy for **G**. In addition, behavioral covariances have been previously estimated in the closely related congener *P. leucopus* (32). All research was conducted in accordance with institutional guidelines (NDSU IACUC A17055) and the guidelines of the Animal Behavior Society (33) and the American Society of Mammalogists (34).

### Trapping and tagging

Individuals for whom phenotypes and selection were estimated were captured using repeated live trapping using Sherman live traps (5.2 × 6.4 × 22.9cm). Traps were set in a 9 × 11 grid, with traps 12.5 m apart, totalling 99 traps. Mixed birdseed and rolled oats were mixed with peanut oil and used to bait traps, and cotton was added to provide insulation. Trapping was conducted between May 30^th^ and October 13^th^ in 2017, with trapping occurring around three times a week. Traps were set between 3-6 pm and checked starting at 6 am the following morning. Closed traps without a capture were counted to allow estimates of trapping effort. Traps with captures were set aside for subsequent behavioural trials and all open traps were closed.

Individual deer mice were tagged with metal ear tags in both ears at first capture for identification at recaptures. All captured individuals were identified by individual ID, had their mass and sex recorded, and identified as either juvenile or adult (developmental stage). Individual ID was recorded to allow subsequent mark-recapture analysis, the estimation of survival, and for repeated behavioral trials.

### Behavioural assays

To investigate the relationships between behavioral syndromes, plasticity, and selection in this population, captured deer mice were tested in three behavioral assays: an open field test, a mirror image stimulation test, and a predator cue response test. All behavioral tests were conducted in arenas (60 × 60 × 40 cm) made of 2 cm thick plywood. Transparent plexiglass was used as lids and a video camera was mounted above each arena (Figure S2). Deer mice were always put through the assays in the same order: 1) open field test, 2) mirror image stimulation test, and 3) predator cue response test. This was done to avoid any carry-over effects from interacting with the mirror on open field tests or from the exposure of the predator cue affecting open field or mirror image response. After the deer mice had been run through all three assays they were released at the same location as they were caught.

#### Open Field Test

In the open field assay, the arena was empty and was used to measure general activity level (35). Activity in an open field arena has been shown to give repeatable measures for general movement in several species of rodents (36, 37) including *Peromyscus maniculatus* (29, 30). At the start of the open field assay deer mice were placed under a cup (11.5 cm in diameter) and given one minute to acclimate. The shelter was then removed and the deer mouse had 6 minutes to explore the arena. We walked at least 20 m away from the recording area to reduce any disturbance caused by our presence. After the deer mouse completed the open field assay, the cup was placed over the deer mouse again, and a cardboard plate was pushed under it to enable us to transport it to the mirror arena. Video analysis tracking was started 2 minutes into the video to make sure that we had time to start the mirror image stimulation and anti-predator tests that were run for other individuals simultaneously (see below) and to move away from the arena area. Distance moved (in cm) in the open field was used as a measure of activity.

#### Mirror Image Stimulation Test

The arena for the mirror image stimulation test had a mirror attached along one wall (Figure S2). Mirror tests have been shown to be an appropriate measure of agonistic behavior in several species of rodents (38-40), and to correlate with aggression towards conspecifics in *D. merriami* (41). Mirror tests also have the advantage of standardizing size differences between the opponents as well as avoiding any injury to individuals being tested. The deer mice were moved from the open field arena into the mirror arena, where they were again given one minute to acclimate under a cup. Video analysis started 1.5 minutes into the trial (again, to avoid any disturbance caused by starting the anti-predator response trial and then move 20m away) and aggression was measured as the time spent on the mirror side of the arena (in seconds). If the deer mice perceived their reflection as a conspecific, spending more time in front of the mirror would indicate interaction with conspecifics, while time spent away from the mirror indicates avoidance.

#### Predator Cue Response Test

The anti-predator response arena had a circular filter paper (11 cm in diameter) soaked in coyote urine placed in the upper left corner (see Figure S2). Small rodents have been shown to have aversive responses towards predator cues (42, 43), including the closely related species *P. gossypinus* (44). Both coyotes (*Lupus latrans*) and red foxes (*Vulpes vulpes*) have been observed in the area, making canid urine generally and coyote urine specifically a suitable cue of predator presence for this deer mouse population. The filter papers were prepared elsewhere and stored at – 20°C until the morning of behavioral trials. After completing the mirror test, a deer mouse was moved to the anti-predator response arena and put under a cup for a 1 minute acclimation period. In subsequent video analysis, tracking started 30 seconds into the video, and anti-predator response was measured as the mean distance from the predator cue (in cm). More responsive individuals were predicted to increase their distance from the predator cue, indicating shy behavior, while bolder and less responsive individuals would stay closer to the cue.

### Statistical analyses

Statistical analyses were conducted using the packages RMark (45) and MCMCglmm (46) in R (47).

### Average Behavioral Responses

Generalized mixed effect models were used to investigate the validity of the behavioral assays by comparing the average responses towards the mirror and the predator cue to behavior in the open field arena using the MCMCglmm package in R. For the mirror assays, the time spent on the mirror side of the arena was compared with time spent on the equivalent side in the open field trial. Similarly, in the anti-predator assay the distance from the predator cue was compared to the distance from the equivalent location in the open field assay. Arena type was added as a fixed effect and animal ID was added as a random effect. Models were run for 650,0000 iterations, with a thinning interval of 5000, and a 150,0000 iteration burn-in. The priors used in all models were minimally informative for (co)variances and flat for correlations. pMCMC values (used similarly to p-values for maximum likelihood statistics) were used to determine whether there was a substantive effect of arena type. pMCMC was calculated here as the proportion of the estimates of fixed effects from the posterior distribution that were below 0. A high proportion (>0.95) indicates that most estimates are below 0, while a low proportion (<0.05) indicates that most estimates are above 0. Both high and low proportion means that the posterior distribution has a very low overlap with 0 and is therefore considered to have a substantive effect on the behaviors.

### Repeatabilities and correlation matrices

Repeatability and among- and within-individual variance-covariance matrices were estimated for distance moved, time spent in front of the mirror, and mean distance from predator cue using multi-response generalized mixed effect models (48, 49). Sex, developmental stage, mass, and the presence of a shelter were included as fixed effects, where mass was within-subject centred (50). Individual ID was included as a random effect. Because distance moved had a left-skewed distribution, this variable was square root transformed to satisfy the assumption of normality prior to all statistical analyses. All three behaviors were mean centered and scaled by their standard deviations to facilitate model fit. The number of iterations, thinning intervals, and burn-in was the same as above. Repeatabilities, correlation matrices, and estimates of individual behaviour (i.e. best linear unbiased predictors, BLUPs) were estimated from the posterior distributions. pMCMC values were used to determine if the fixed effects had any substantial effect on the behaviors (see above).

### Selection gradients

Directional selection gradients (**β**) for the three behaviors (activity, aggression, and anti-predator response) were estimated from the effect of individual behavior (as BLUPs from the multi-response generalized mixed effect model) on survival probability from mark-recapture models fitted using Program MARK (45) and RMark. Sex, developmental stage, mass, and the BLUPS for distance moved (activity), time spent in front of the mirror (aggression), and mean distance from predator cue (anti-predator response) were included as covariates. We assumed a closed population and recapture probability was set as constant throughout the season. Because some of the individuals were too young to identify sex, sex was recorded as female (1 0), male (0 1), or unidentified (1 1). The coefficients for survival for the three behaviors were converted to a traditional Lande and Arnold (10) directional selection gradient (**β**) using the method detailed in Waller and Svensson (51).

To investigate whether behaviors had a substantive effect on survival, seven additional models were fit to allow for comparison (Table 1). If any of the behaviors had a substantial effect on survival, models 1 – 7 would have a better fit than the model with no behavior covariates (model 8). Relative model fit was determined based on AICc values

**Table 1.** The eight mark-recapture models fit to investigate the effect of behavior on survival.

| Model | Behavioral covariates |
| --- | --- |
| 1 Effect of all behaviors on survival | Activity, aggression, and anti-predator response |
| 2 Effect of activity and aggression on survival | Activity and aggression |
| 3 Effect of activity and anti-predator response on survival | Activity and anti-predator response |
| 4 Effect of aggression and anti-predator response on survival | Aggression and anti-predator response |
| 5 Effect of Activity on survival | Activity |
| 6 Effect of aggression on survival | Aggression |
| 7 Effect of anti-predator response on survival | Anti-predator response |
| 8 No effect of behaviors on survival | No behaviors |

### Alignment

We used the among- and within individual covariance matrices (**I-** and **W-**matrices) as proxies for genetic covariation and multivariate expression of adaptive plasticity, respectively, to test predictions regarding alignment. **i_max_** is the dominant eigenvector for the **I**-matrix and is the dimension with the most among-individual variation while **w_max_** is the same for the W-matrix. The dominant eigenvectors for both the among- and within-individual covariance matrices were estimated from their posterior distributions estimated by the multi-response mixed effects model (1000 estimates in total). These were then used to calculate the vector correlation and the angle between (i) **i_max_** and **w_max_**, (ii) **i_max_** and the selection gradient (**β**), and (iii) **w_max_** and **β**. Vector (V) correlations were calculated as:

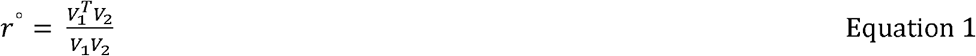

where V_1_ and V_2_ represent **i_max_**, **w_max_**, or **β**. The vector correlation was then used to estimate the angle between vectors:

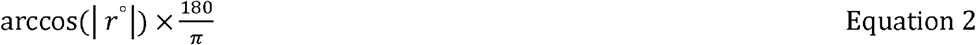

A vector correlation that does not differ from 1 (i.e. an angle that does not differ from 0) would mean that the vectors are aligned.

Because we were interested in whether a vector correlation was statistically distinguishable from 1, rather than common null expectation of 0, standard statistical analyses were not appropriate. We therefore used a Bayesian approach developed by Ovaskainen, Cano (52), which uses posterior distributions. Using the posterior distribution from the multi-response mixed effects model, we obtained 1000 estimates of the dominant eigenvectors for the among- and within-individual correlation matrices. To get similar estimates for the selection gradients, the full mark-recapture model described in Table 1 (Model 1) was re-fit using the 1000 posterior estimates of the BLUPs for the three behaviors from the multi-response mixed effects model. Probability of alignment between among- and within-individual eigenvectors, among-individual eigenvectors and selection gradients, and within-individual eigenvectors and selection gradients was then estimated as (5, 52):

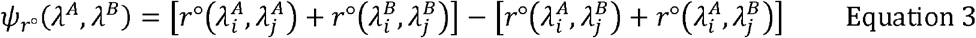

where ⍰^A^ and ⍰^B^ represents the posterior distributions of vectors (**i_max_**, **w_max_**), or the 1000 estimates of **β**), and i and j represents different samples from these distributions. The correlation between samples of the same vectors should be 1, thus the first part of the equation = 2 in the absence of estimation uncertainty. The inclusion of this term incorporates estimation uncertainty present in the posterior distribution. The second part of the equation is the correlation between estimates of different vectors. If two vectors are highly correlated, the second part of the equation is also ≈ 2, and ⍰_r°_ ≈ 0, with sampling error allowing ⍰_r°_ to be below zero. As the correlation between ⍰^A^ and ⍰^B^ decreases, ⍰_r°_ approaches 2. If 95 % of the estimated ⍰_r°_ are above 0, the two matrices are considered misaligned. We therefore calculated the proportion of ⍰_r°_ values that were positive, with a higher value meaning that more estimates exclude 0, and will in this case indicate a substantial misalignment.

Unfortunately the method of Ovaskainen, Cano (52) can be sensitive to low power so to further understand the alignment between vectors we also compared the estimated vector correlation to three different null expectations of the correlation following Berdal and Dochtermann (5). The null expectations where set to 0.975, 0.95, and 0.9, which are considered highly correlated and indicates that the vectors are aligned. The differences between observed and null correlations were then transformed to z-values. Z-values larger than 0 would indicate that the estimated vector correlation was higher than the null expectation, which would mean that the vectors were aligned. 1000 estimates of vector correlations between **i_max_** and **w_max_**, between **i_max_** and **β**, and between **w_max_** and **β** were compared to the null expectations. We then calculated how many of the Z-values were above 0, i.e. how many of the estimated vector correlations were above null expectations. As before, if more than 95% of the estimates are above this value the misalignment was considered to be substantial.

### Testing for historical and stabilizing selection

After comparing the eight mark-recapture models (Table 1) we found that the model without behavioral traits had the best fit, indicating that the behaviors measured here were under weak or no directional selection. Two post-hoc analyses were therefore carried out to investigating variability and the possibility of stabilizing selection for the three behavioral traits (see Discussion), as both low variation and stabilizing selection will reduce alignment with the behavioral variation and selection. The first post-hoc analysis was the calculation of mean standardized variances for the three behaviors, i.e. evolvability, I_A,_ (53).

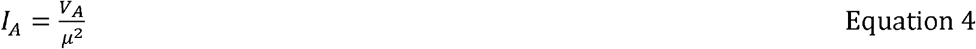

where V_A_ is the additive genetic variance of a trait, and μ is the mean of the same trait. This gives an estimate of the evolutionary potential of a population. Mean-scaling the variances allows us to compare these estimates across traits and species. Low values of evolvability suggest that a trait has been under strong historical selection, leading to a depletion of variation. However, in our case evolvability is estimated based on among-individual variance instead of just the additive genetic variance, meaning our estimate also included dominance- and epistatic variance as well as permanent environmental effects (54). The estimated intercepts from the multi-response generalized mixed effect models were used for the behavioral means in Equation 4.

Second, we added quadratic terms for behaviors to the mark-recapture model (10) where a negative coefficient on the quadratic term would indicate stabilizing selection (55). As before, eight models were fit (Table S1) and AICc values were used to compare the model fit. All models included sex, developmental stage, mass, and the linear terms (i.e. the BLUPS) for the behaviors, and only differed in the number of quadratic terms for the behaviors that were added (none, one, two, or three quadratic terms). If the behavioral traits were under stabilizing selection, models including these terms and with negative coefficients should have a better fit.

## Results

We sampled the population for 30 nights and captured a minimum of 92 individuals (including individuals that escaped prior to being tagged or identified) with a mean recapture rate of 3. Of these, 72 individuals were behaviorally assayed and used in analyses. This included 32 females, 36 males, and 4 individuals of unidentified sex, where 48 were adults and 24 were juveniles. The other 20 individuals managed to escape either while being tagged or during the trials and were never caught again, meaning they had insufficient behavioral data collected to be used in our analyses. In total, we conducted 641 behavioral assays (Table ***2***), with a mean number of trials per individual of 3.13±2.65, 2.65±2.23, and 2.79±2.56, for activity, aggression and antipredator response respectively.

**Table 2.**
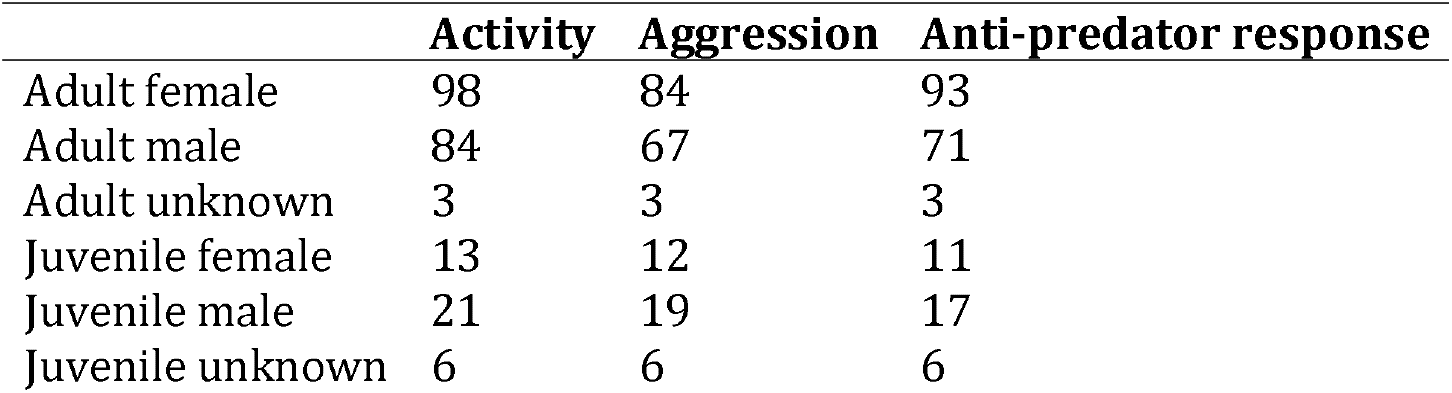
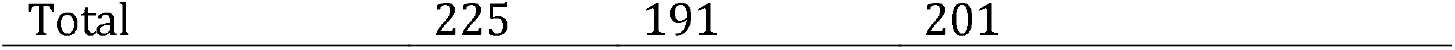
Number of trials for the different sexes and developmental stages.

|  | Activity | Aggression | Anti-predator response |
| --- | --- | --- | --- |
| Adult female | 98 | 84 | 93 |
| Adult male | 84 | 67 | 71 |
| Adult unknown | 3 | 3 | 3 |
| Juvenile female | 13 | 12 | 11 |
| Juvenile male | 21 | 19 | 17 |
| Juvenile unknown | 6 | 6 | 6 |

### Average Behavioral Responses

We found that deer mice responded as predicted to the predator cue, increasing their distance from the cue in the predator response test compared to the corresponding corner during the open field test (pMCMC = 0.00, Figure S3a). Deer mice also spent more time on the mirror side of the arena compared to the controlled open field test (pMCMC = 0.008, Figure S3b). In addition, repeatability for staying on the mirror side of the arena was 0.36 for the mirror-trial, while for the open field trial side preference was indistinguishable from 0, indicating that the deer mice stayed on one side of the arena consistently when the mirror was present but not when absent, providing further validation of the assay.

From the multi-response generalized mixed effect model we found that males had a higher activity level compared to females, 0.47 (0.08 – 0.89), pMCMC = 0.007, and the presence of a shelter reduced the distance moved in the open field, −1.04 (−1.44 – −0.67), pMCMC = 1, and anti-predator assay, −0.41 (−0.87 – 0.03), pMCMC = 0.96 (Table S2). No other fixed effect substantively affected the assayed behaviors (Table S2).

### Behavioral Variation

All behaviors were moderately repeatable: activity, aggression, and anti-predator response had estimated repeatabilities of 0.47 (0.35-0.60), 0.36 (0.24-0.50), and 0.33 (0.21 – 0.46) respectively (Figure 1).

**Figure 1.**
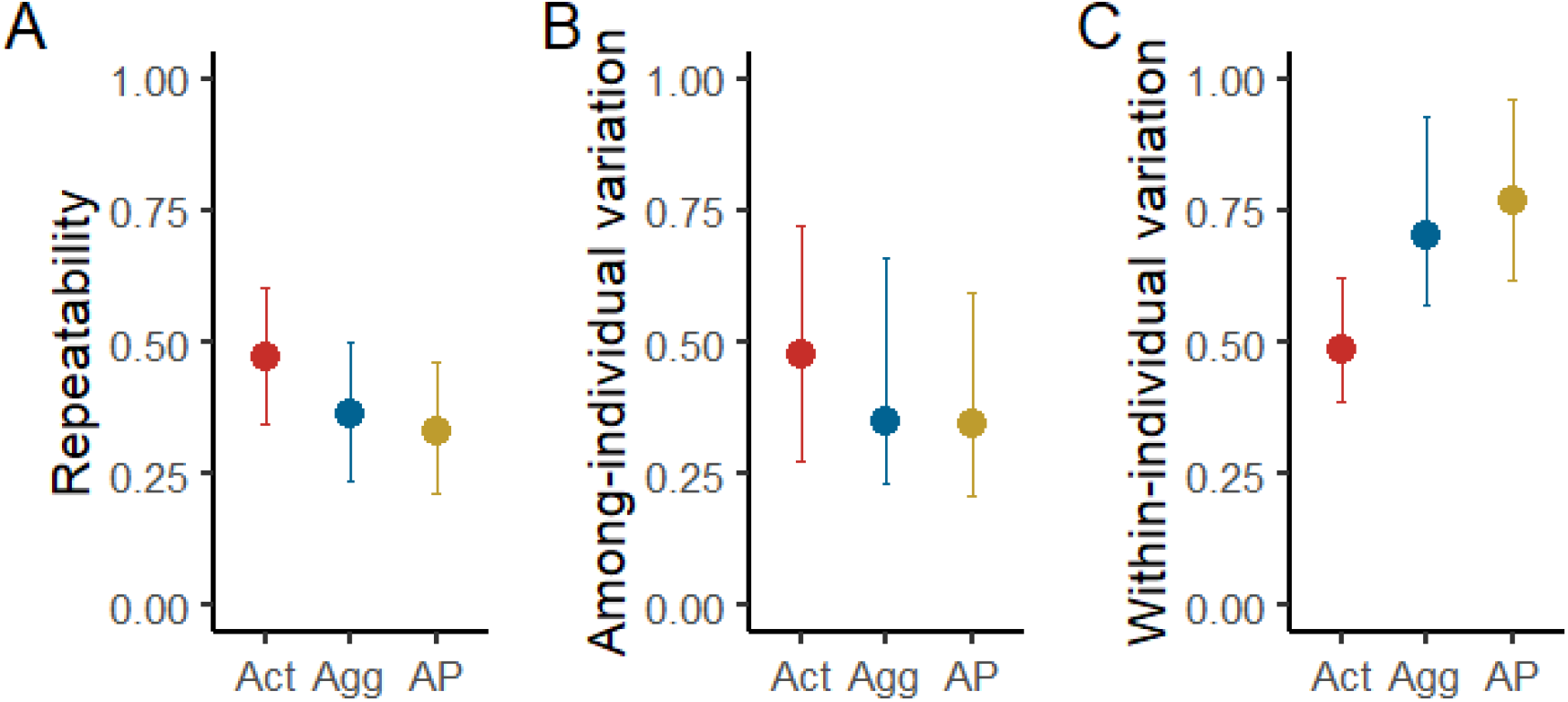
The A) repeatability, B) among-individual variation, and C) within-individual variation for activity level (red), aggression (blue), and anti-predator response (yellow).

### Correlation matrices

Anti-predator response and activity in the open field were negatively correlated at both the among- and within-individual levels. Put another way, less active individuals stayed further away from the predator cue and individuals that reduced their activity level also increased their distance from the predator cue (Table **3**). The among-individual correlation between predator cue response and aggression was around 0, while the within-individual correlation was slightly negative (Table **3**). This negative correlation suggests that when individuals increased their time in front of the mirror, they also reduced their distance to the predator cue. The largest difference between the among- and within-individual correlation was for the relationship between activity and aggression (Table **3**). Here, the among-individual correlation was negative while the within-individual correlation was positive. This means that while individuals that are more active spend less time in front of the mirror, an individual that increases its activity level will also increase its time spent in front of the mirror. However, only the within-individual correlations between activity and aggression and between activity and anti-predator response were substantively different and statistically distinguishable from zero (Table **3**).

**Table 3.** Among-individual correlations (above the diagonal) and within-individual correlations (below the diagonal) between activity (Act), aggression (Agg), and anti-predator response (AP). Substantive correlations (pMCMC > 0.95 or pMCMC < 0.05) are bolded. Evolvabilities are shown on the diagonal.

|  | **Act** | **Agg** | **AP** |
| --- | --- | --- | --- |
| **Act** | 0.22 (0.08 – 0.36) | -0.25 (-0.54 – 0.07)<br>pMCMC = 0.92 | -0.25 (-0.59 – 0.05)<br>pMCMC = 0.91 |
| **Agg** | **0.17 (0.03 – 0.36)**<br>pMCMC = **0.01** | 0.03 (0.003 – 0.08) | -0.06 (-0.32 – 0.40)<br>pMCMC = 0.50 |
| **AP** | **-0.14 (-0.31 – 0.02)**<br>pMCMC = **0.96** | -0.16 (-0.30 – 0.04)<br>pMCMC = 0.94 | 0.01 (0.001 – 0.03) |

### Mark-Recapture Results

Only mass had a substantive effect on survival and only developmental stage had a substantive effect on recapture probability (Table S3). Larger individuals were found to have a higher survival probability than smaller individuals and adult individuals had a higher chance of being recaptured compared to juveniles. The selection coefficient for the behaviors were relatively low, especially for aggression. This low effect of behaviours on survival is further emphasised by the model comparison results, where the model with no behavioral terms had the lowest AICc score (Table S4). The same results were found when investigating the presence of stabilizing selection on the behaviors, where the model with no quadratic terms had the best fit (Table S5, Table S6). However, the model without behaviors did not statistically differ from one including a quadratic term for aggression (AICc < 2 points away from the reduced model), indicating the possibility stabilizing selection on aggressive behavior.

Despite the lack of clear effects on survival, we converted the coefficients for the effects of behaviors on survival to Lande and Arnold’s directional selection gradients (**β**, Lande and Arnold, 1983) which were used to estimate vector correlations and angles with among-and within-individual eigen vectors. The **β**s were −0.67 (−1.23 – 0.45), 0.23 (−1.10– 1.19), and −0.62 (−1.22 – 1.19) for activity, aggression, and anti-predator response, respectively.

### Vector correlations and angles

The vector correlations between the dominant eigenvectors for the among- and within-individual covariance matrices, and with selection gradients were low, leading to angles around 70° to 90° (Figure 2, Table S7).

**Figure 2.**
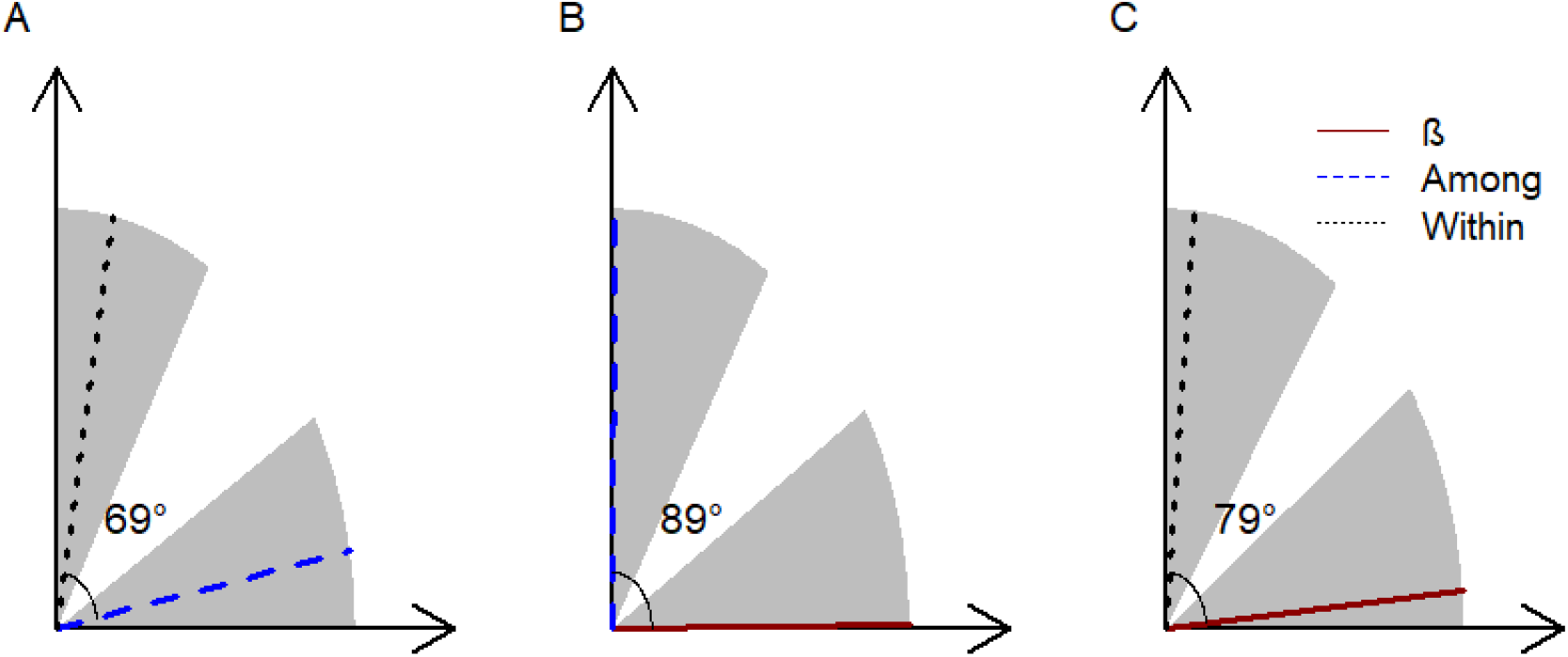
Angles between A) **i_max_** (dashed blue line)and **w_max_** (dotted black line), B) **i_max_** and **β** (solid red line), and C) **w_max_** and **β**. The angles between lines are based on the posterior distribution from the multi-response mixed effects model, and the shaded areas show the 95% HPD interval, where the non-shaded area indicates the minimum HPD interval.

Using the Bayesian method for matrix comparison described by Ovaskainen, Cano (52), the three vectors were not clearly misaligned, but the probability of misalignment (i.e. proportion of estimates > 0) was quite high, particularly between **i_max_** and **w_max_** (Table 4). Because this Bayesian method is sensitive to low power, we also investigated the alignment using Z-transformation. This alternative method indicated that most of the estimated vector correlations were less than 0.9 (Table 4), consistent with genetic variation, plasticity, and selection gradients being misaligned.

**Table 4.**
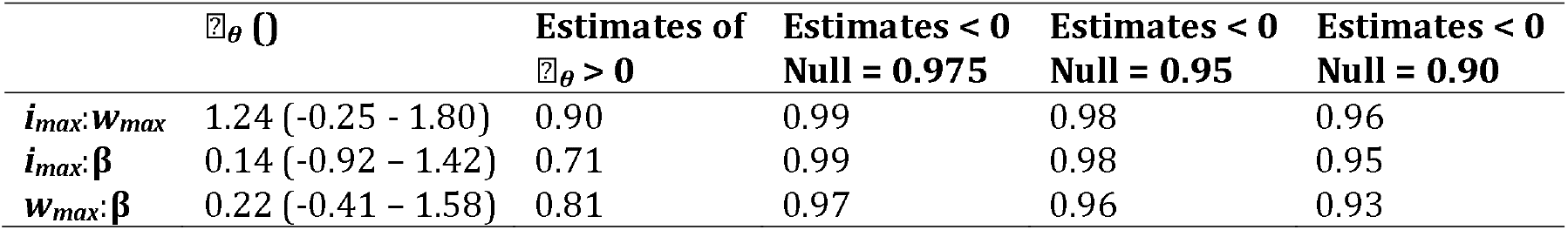
⍰_θ_ and proportion of ⍰_θ_ > 0 for the alignment between among – and within-individual variation (**i_max_** and **w_max_** respectively) and selection gradient (***β***), and proportion of estimates (out of 1000) that are below three different null expectations (correlation of 0.975. 0.95, and 0.90) for the vector correlation between **i_max_**, **w_max_**, and **β**. Because the proportion of ⍰θ estimates > 0 was less than 0.95 none of the vectors were clearly misaligned. However, the vector correlations where substantially different from the three null expectations (except for **w_max_**:***β*** compared to the null expectation of a correlation of 0.90), indicating a misalignment between the three vectors.

| | $\mathbb{Z}_{\theta}$ | Estimates of $\mathbb{Z}_{\theta} > 0$ | Estimates $< 0$<br>Null = 0.975 | Estimates $< 0$<br>Null = 0.95 | Estimates $< 0$<br>Null = 0.90 |
| --- | --- | --- | --- | --- | --- |
| $i_{\max}:w_{\max}$ | 1.24 (-0.25 - 1.80) | 0.90 | 0.99 | 0.98 | 0.96 |
| $i_{\max}:\beta$ | 0.14 (-0.92 - 1.42) | 0.71 | 0.99 | 0.98 | 0.95 |
| $w_{\max}:\beta$ | 0.22 (-0.41 - 1.58) | 0.81 | 0.97 | 0.96 | 0.93 |

## Discussion

Contrary to our predictions we found little evidence for alignment between genetic variation (among-individual variation), plasticity (within-individual variation), and selection gradient in the wild population of deer mice. Most estimates of alignment were greater than 0 (0.71– 0.90, Table 4), indicating that most of the vectors are different from each other. In addition, the results from the Z-transformation analysis indicated that all the three vectors have a vector correlation substantially different from 1, i.e. they are misaligned (Table 4). A misalignment between among-individual variation and the selection gradient is predicted when there has been a change in selection pressure and the genetic architecture has not had enough time to realign with the fitness landscape (8). However, because phenotypic plasticity can respond quicker to selection pressure (16), plasticity is still predicted to align with selection. This was not the case in our study system, and there are at least three reasons for the apparent misalignment between the selection gradient and both among- and within-individual variation.

First, the estimates of the selection gradient might be biased. Developmental stage had an effect on recapture probability, with adult individuals having a higher chance of being recaptured than juveniles. Juvenile deer mice have a higher rate of dispersal at the end of the breeding season (56), which might be why recapture probability was lower for juveniles in our system. One of the assumptions of the mark-recapture model used is that we were monitoring a closed population, meaning that an individual that has not been recaptured for a long time is assumed to be dead. However, this was not necessarily the case for our population, and deer mice were able to migrate out of our trapping location, which means that there might be some biases in the survival estimates. Unfortunately, given the structure of our data, this assumption was necessary to estimate the effect of behaviors on survival.

Second, the behaviors measured here might be under weak or no directional selection. Consistent with this, only mass had a substantive effect on survival probability, with larger individuals having a greater chance of surviving compared to smaller individuals. Including behavior in the mark-recapture model did not improve the model’s fit (Table S4). This indicates that the behavioral variables had only small (or no) effects on survival probability, and that they are not under strong directional selection. If the selection pressure is weak it might not be strong enough to have shaped **G** to align with selection. Thus, the behaviors measured here might not have been appropriate for this population of deer mice. However, the deer mice increased their time spent on the mirror side of the arena in the mirror image stimulation test compared to the open field test (Figure S3a), and likewise increased their distance from the predator cue compared to the same corner in the open field test (Figure S3b). This indicates that the deer mice did respond to these cues in the predicted manner and validates the use of these behaviors in this study. All behaviours were also repeatable, meaning that deer mice were consistent in their responses to these cues as well as in their activity level. In addition, previous studies have shown that there is additive genetic variation for distance moved (31) and underlying genes influencing aggressive behavior (57) for *P. maniculatus,* further supporting the suitability of these methods to record the behaviors used in this study.

While these behaviors not being under selection is perhaps the most parsimonious explanation for our result, such a finding would be particularly surprising. The behaviors we measured are frequently associated with fitness across taxa (58). Exploration, response to cues of predator presence, and mirror image stimulation are also frequently argued to be particularly ecologically meaningful (59-62). For example, rats (*Rattus spp*.) decrease their foraging time in the presence of a fresh predator cue (63), and female red squirrels (Tamiasciurus hudsonicus) with a higher activity level were more likely to be risk-prone which lead to a lower winter survival for the female herself, but a higher survival of her offspring because the offspring could remain in their natal territory (39).

Third, the behaviors might be under stabilizing selection, or be at their fitness peak, or have low genetic variation. Only traits under directional selection are predicted to align with the selection gradient (5). If the behaviors are under stabilizing selection, the variation of the traits will be around the fitness optima and variation in the traits is predicted to be orthogonal to selection. Consistent with this Blows, Chenoweth (64) found that the genetic correlations among cuticular hydrocarbons in male *Drosophila serrata* were misaligned with directional selection, most likely due to a reduction in genetic variation caused by sexual selection through strong female preference. Thus, both weak directional selection and stabilizing selection could explain the misalignment of genetic variation (here among-individual variation) and the selection gradient.

However, we could not determine whether the behaviors were under stabilizing selection based on our original a *priori analysis*. We therefore carried out the described post-hoc analysis to estimate stabilizing selection by adding quadratic terms to the mark-recapture model (10). These results were similar to the analysis of linear terms, and the behaviors were not clearly under stabilizing selection (Table S5, Table S6). Specifically, the model without any quadratic terms for behaviors had the best fit but was not distinguishable from the model including a quadratic term for aggression. The coefficient for aggression in this was negative (Table **3**), indicating that aggression could be under stabilizing selection.

In contrast, aggression has been found to be under stabilizing selection in both water voles (*Arvicola terrestris*) and house mice (*Mus musculus*) as a result of female choice, where females favoured males with a medium aggression level over overaggressive or docile males (65). Similarly, female meadow voles (*Microtus pennsylvanicus*) avoid mating with overaggressive males (66), thus hindering selection for increased aggressiveness in male voles, i.e. aggression is not under directional selection. Sexual selection on aggression could also be the case for our population of deer mice and could reduce the alignment between among-individual variation and selection. Unfortunately, at this time, stabilizing selection cannot be clearly distinguished from no selection.

Haller and Hendry (67) demonstrated that selection in traits that have reach their fitness peak is difficult to detect and we therefore sought to explore the possibility of stabilizing selection further. Another potential indicator of populations being under strong selection is a loss of variation (Mousseau and Roff 1989, Houle 1992). We therefore calculated the mean standardized among-individual variation of the behaviors (i.e. evolvability, Ii). These values were low compared to the estimates in Hansen, Pélabon (53), indicating low evolvability in these behavioral traits (Table **3**). This observed low I_i_ is consistent with behaviors being close to the fitness peak for this population, which would also explain the misalignment of among- and within-individual variation with the selection gradient.

As mentioned, among- and within-individual covariation were also misaligned. The among- and within-individual correlations were low to moderate, and only the within-individual correlation between activity and aggression and activity and anti-predator response were found to be substantive (pMCMC > 0.95 or < 0.05, Table **3**). The lack of substantial correlation for the among-individual behaviors is surprising, as this has been found in several other rodents (68, 69), including closely related species (32).

The within-individual correlation between activity and anti-predator response were negatively, meaning individuals that reduced their activity level from one trial to another also increased their distance from the cue. Activity and aggression, on the other hand, were positively correlated at the within-individual level, which means that individuals that increase their activity level would also increase their aggression towards a conspecific. However, the among-individual correlation between activity and aggression was negative, indicating that individuals with higher activity levels would be less aggressive. The opposite signs for the correlations between activity and aggression might be the main reason why we did not observe an alignment between the among- and within-individual covariance matrices. Within-individual variation includes other forms of phenotypic plasticity in addition to adaptive plasticity (5, 27, 28), and only adaptive plasticity is predicted to align with the selection surface and genetic variation, whilst other sources of plastic variation (e.g. passive plasticity, measurement error, and organismal error) are not (5). Thus, the proportion of the within-individual variation that is made up by adaptive plasticity determines how well within-individual variation should align with both genetic variation and the selection surface (see worked example in 5). Here, our finding of misalignment also shows the importance of separating among- and within-individual covariation as they can differ greatly, and because they are produced by differently and have different implications (5, 49, 70).

There are several models examining how genetic and plastic variation is shaped by selection as well as how they affect each other (7, 16, 71, 72), but few empirical studies have addressed this topic (but see 73). Here we have shown that the relationship between a selection gradient and among- and within-individual covariation in a wild population can be estimated using Bayesian methods. Overall, we found no evidence for alignment among these three characterizations of phenotypes and selection in this wild population of deer mice. The lack of alignment combined with weak or no selection on these behaviors and the low evolvabilities suggests the possibility that the behaviours measured here might be close to their fitness optimum. Furthermore, the low evolvabilities indicate that this population has very low potential of responding to a change in selection pressure. Subjecting a sample of this population to a new selection regime in a lab setting would give us a better idea of the adaptive potential of this population, and whether there is enough phenotypic plasticity to overcome the apparent lack of genetic variation suggested by the low evolvability of each trait.

## Acknowledgements

This study was funded by Animal Behavior Society (ABS Student Research Grant, 2017), the National Science Foundation (US NSF IOS 1557951), and the Biology Department at North Dakota State University. We thank Erin Mollberg, Linda Rausch, and Elizabeth Kieffer for helping with the data collection. We also thank Steven Travers, Julia Bowsher, Lauren Hanna, Courtney Hickey, Jeremy Dalos, and Julian Evans for helpful discussion. Special thanks to Raphaël Royauté for statistical help as well as feedback on earlier drafts.

